# Nanobody-mediated modulation of long RSH enzymes Rel and RelA catalysis by restriction of their conformational landscape

**DOI:** 10.1101/2025.08.07.669093

**Authors:** Katleen Van Nerom, Andres Ainelo, Kyo Coppieters ’t Wallant, Ariel Talavera-Perez, Dannele Echemendia-Blanco, Sarah Peeters, Brahim El Khalfaoui Oulali, Hedvig Tamman, Tatsuaki Kurata, Mohammad Roghanian, Chloé Martens, Els Pardon, Jan Steyaert, Vasili Hauryliuk, Abel Garcia-Pino

## Abstract

Long RSH enzymes, Rel and RelA, are the master regulators of (p)ppGpp alarmone levels in bacteria. Their catalytic activity is governed by transitions between a compact, hydrolysis-competent (HD^ON^) state and an elongated, synthesis-competent (SYNTH^ON^) state. The equilibrium between these states is modulated by factors such as “starved” ribosomes and regulatory proteins DarB, EIIA^NTR^, ACP and YtfK. Here, we identify and characterize camelid nanobodies that act as selective allosteric modulators by trapping Rel/RelA enzymes in distinct conformational states. Nanobodies that lock the TGS domain of RelA and prevent its activation by deacylated tRNA on starved ribosomes, strongly inhibit (p)ppGpp synthesis and suppress the virulence of *E. coli* in an animal model. Nb898 stabilizes Rel in the open SYNTH^ON^ state, enhancing synthesis activity while suppressing hydrolysis. Conversely, Nb585 traps Rel in a HD^ON^ conformation, strongly inhibiting alarmone synthesis while promoting (p)ppGpp hydrolysis. Structural and biochemical analyses reveal that nanobodies, like natural allosteric regulators, act by restricting the RSH enzyme’s conformational landscape. These findings establish nanobodies as powerful tools for dissecting RSH function and provide potential leads for developing protein-based RSH modulators.

## Introduction

Nucleotides guanosine-3’,5’-tetraphosphate and guanosine-3’5’-pentaphosphate – collectively referred to as (p)ppGpp alarmones – play a key role in bacterial stress sensing and adaptation^1,2^. The alarmones control different aspects of physiology including the growth rate, metabolism, transcription and translation and are effective at different cellular concentrations^3–8^. As master regulators of bacteria physiology, the alarmones are also implicated in control of bacterial virulence, biofilm formation and tolerance to antimicrobials^9–13^.

The balance between synthesis and hydrolysis of (p)ppGpp is tightly controlled. The primary enzymes controlling (p)ppGpp homeostasis are “long” Rel/RelA/SpoT homolog enzymes, collectively known as long RSHs^14,15^. These enzymes are near-universal in bacteria, present in genomes of the majority of species, with an exception of Plantomycetes, Verrucomicrobia and Chlamydiales, the PVC superphylum^14^. Long RSHs contain two catalytic or pseudo-catalytic domains in their N-terminal half (NTD): a hydrolysis domain (HD; or pseudo-HD if inactive) and a synthesis domain (SYNTH; or pseudo-SYNTH if inactive). Their enzymatic activity is regulated by four C-terminal regulatory domains (CTD)^16–20^. In the case of Rel and RelA, the primary signal that stimulates (p)ppGpp production is amino acid starvation, which is sensed by recognizing the deacylated (uncharged) tRNA in the ribosomal A site^21,22^. In addition to this *in cis* intra-protein regulation, the SYNTH activity of Rel/RelA is also stimulated *in trans* by the alarmones themselves, with pppGpp being a more potent activator^17,23,24^.

A complex allosteric network controls and finetunes the enzymatic output of long RSH enzymes. In the case of bifunctional (capable of both degrading and synthesizing the alarmones) RSH enzyme Rel the opposing activities of both catalytic domains are controlled by the relative concentrations of substrates, with the ligand binding to either of the active sites switching off the activity of the other^19^. Importantly, the allosteric effect of the CTD supersedes the crosstalk between catalytic NTD domains^17,25^. In the HD^ON^ state, the enzyme is in a closed conformation, in which the CTD activates hydrolysis by stabilizing the HD domain and promoting the organization of the active site while precluding the activation of the SYNTH domain^18^. Upon amino acid starvation, binding of the enzyme to “starved” (i.e. containing a deacylated tRNA in the A site) ribosome in a fully extended elongated state abolishes the allosteric effect of the CTD on hydrolysis and exposes the SYNTH domain^26,27^. However, the complete activation of the SYNTH domain also requires the binding of alarmones to a dedicated allosteric site that, acting in concert with the A-site-bound uncharged tRNA, stabilizes an elongated ribosome-bound state of the enzyme, triggering the processive synthesis of (p)ppGpp^17,25^. By contrast, active hydrolysis is contingent on a compact τ-state of long RSH enzymes Rel and SpoT, in which the SYNTH active site is occluded and the HD catalytic residues and metal co-factor are stabilized and primed for (p)ppGpp binding and hydrolysis^18,19^.

In addition to being controlled by starved ribosomal complexes, NTD and CTD regions are the target of diverse regulatory proteins the modulate the activity of RSHs^28–34^. While EIIA^NTR^, ACP and YtfK were reported to inhibit the hydrolase function of Rel and SpoT^31,34,35^ and the synthetase activity of RelA^36^, DarB activates the synthetase activity of Rel in a ribosome-independent manner^37^. This suggests that long RSH enzymes possess a network of allosteric “hotspots” that modulate the enzymes’ catalytic activities. Such hotspots would allow long RSH enzymes to sense the bacterial metabolic state while simultaneously providing the checkpoints that are crucial to control overproduction of (p)ppGpp^17,18^.

Camelid nanobodies are well stablished structural biology tools used to study the conformational landscape of proteins^38–40^. Here, we leveraged nanobodies to investigate the conformational dynamics of Rel and RelA, identifying allosteric hotspots that can be targeted to modulate their catalytic activity. Our structural and biochemical analyses reveal that by locking Rel/RelA enzymes in specific conformational states, we can precisely manipulate its activity. We have identified and characterized high affinity nanobodies that can mimic the effect of 70S ribosomes on Rel’s synthetase activity by binding to an allosteric site on the SYNTH domain. In contrast, nanobodies that interact with the catalytic domains promote hydrolysis and strongly inhibit synthesis by stabilizing the enzyme’s τ-state. These findings highlight the potential of targeting allosteric sites outside the active centers of RSH enzymes, paving the way for novel antimicrobial strategies against stringent response regulators.

## Results

### Generation of high affinity nanobodies against RSH enzymes

While the structures of the extended conformation of the ribosome-bound state of the (p)ppGpp synthetases Rel and RelA are well characterized^26,27^, their off-ribosome state remains refringent to structural biology. Only the structure of the monofunctional hydrolase SpoT from *Acinetobacter baumannii* was recently determined^18^, which a revealed a compact τ-shaped architecture incompatible with alarmone synthesis. To generate the nanobodies we could use to explore the conformational landscape of the long RSH synthetases off the ribosome, we immunized different llamas with *Escherichia coli* RelA (RelA*_Ec_*) and Rel from *Chlorobaculum tepidum* (Rel*_Ct_*), and obtained camelid nanobodies after phage display selection using established protocols^41^. After two rounds of selection against RelA*_Ec_* and Rel*_Ct_* – either in unbound state or in the presence of GDP or APCPP – we have successfully isolated a set of different candidate binders that were classified according to the sequences of the third complementarity determining region (CDR3, see **Supplementary Fig. 1a**).

### RelA_*Ec*_-targeting nanobodies inhibit (p)ppGpp synthesis by *E. coli* RelA *in vivo*

(p)ppGpp alarmones are essential for maintaining the cellular homeostasis, specifically for controlling the amino acid synthesis^15,42^. This key role of (p)ppGpp is leveraged in so-called SMG functional test for RelA SYNTH activity: only *E. coli* expressing SYNTH-competent RSH can grow in the SMG defined medium^43^. Therefore, to characterize the anti-RelA*_Ec_* nanobodies, we have first expressed them in *E. coli* and monitored the effect on bacterial growth either under permissive conditions (M63 medium^44^, under these conditions inhibition of RelA SYNTH activity does not affect the growth) or under non-permissive conditions (SMG medium, under these conditions inhibition of RelA SYNTH activity abrogated growth).

None of nanobodies were toxic when expressed in bacteria growing on M63 media (**Fig. 1a**). Conversely, IPTG-inducible expression of Nb94 fully abrogated the *E. coli* DJ624 growth on SMG medium but not in M63 (**Fig. 1a**). Nb96 displayed a mild SMG-specific inhibitory effect and Nb120, a control nanobody selected against a different target, did not inhibit growth on SMG. Next, we assessed the interactions between nanobodies and full-length *E. coli* RelA (RelA*_Ec_*) *in vitro* using Isothermal Titration Calorimetry (ITC). Nb94 and Nb96 bound RelA*_Ec_* with affinities of 13 nM and 157 nM, respectively, and no interaction was detectable for Nb120 (**Fig. 1b-d** and **Supplementary Table S1**). Thus, the high affinity of Nb94 and Nb96 to RelA were consistent with the SMG assays that showed these are potent inhibitors of RelA’s SYNTH activity *in vivo*.

**Fig. 1.**
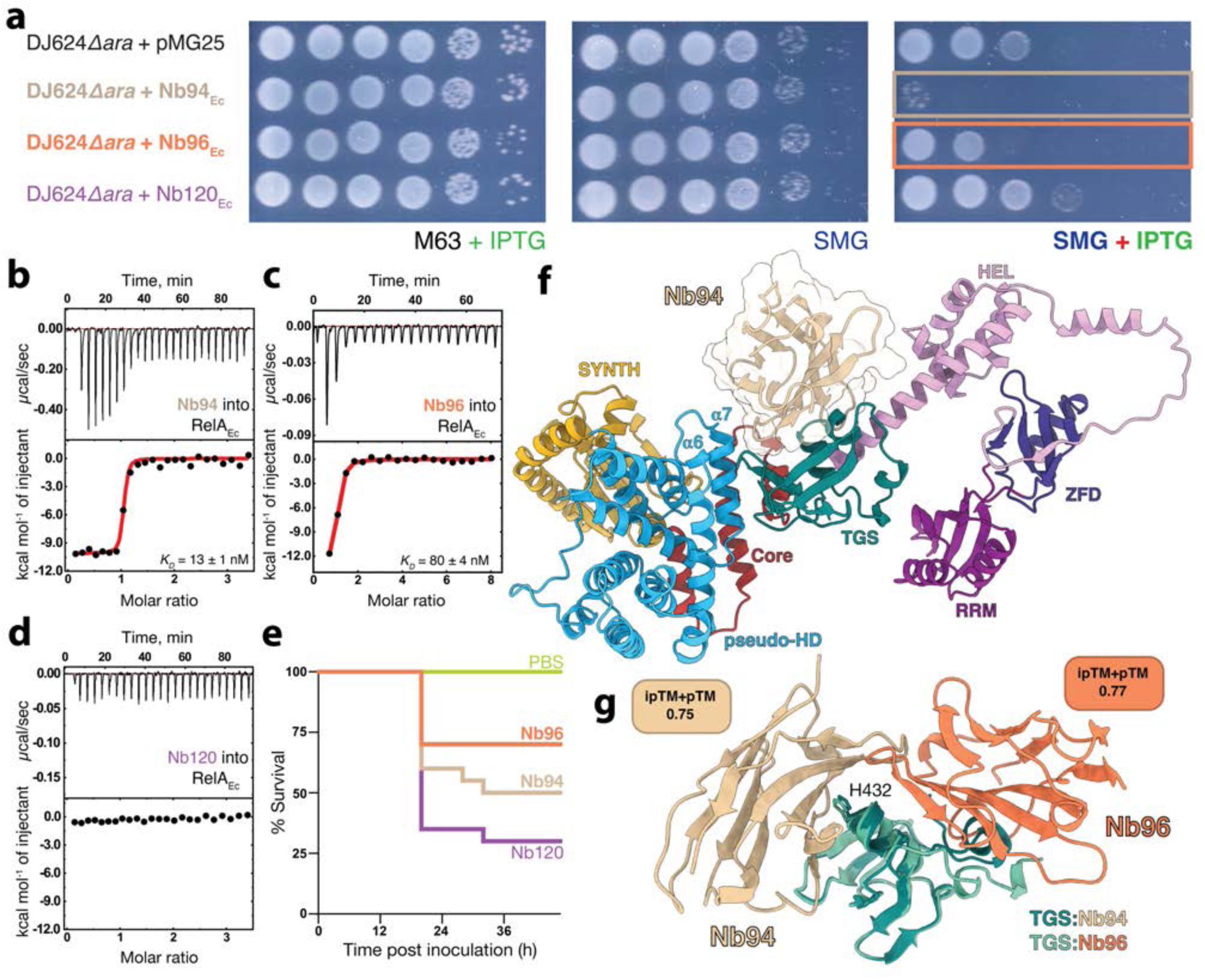
Nanobodies against RelA prevent growth of *E. coli* under amino acid starvation conditions and curtail virulence. **(a)** SMG RelA functionality test of cells transformed with vectors expressing Nb94, Nb96 and the control Nb120 used as negative control. ITC titrations of Nb94 **(b)**, Nb96 **(c)**, and Nb120 **(d)** into RelA. **(e)** Virulence assays in the *G. mellonella* infection model demonstrate nanobodies that inhibit RelA affect strongly the virulence of *E. coli*. **(f)** AlphaFold predicted structural model of the complex of RelA; coloured from N to C-terminus: NTDs pseudo-HD (light blue), SYNTH (in gold) and Core (in red), and CTDs, TGS (teal), HEL (pink), ZFD (dark navy blue) and RRM (purple) and Nb94 (in kaki). **(g)** Details of the interaction of Nb94 (kaki) and Nb96 (orange) with RelA’s TGS domain (coloured in teal and turquoise for each respective complex). The orientation of the nanobodies in both complexes underline the importance of H432 (labeled in the figure) to the binding interface. Prediction scores are shown in the figure.

### Nb94 and Nb96 curtail virulence of *E. coli*

(p)ppGpp-mediated signaling is crucial in antibiotic tolerance and virulence of *E. coli*. We reason that our RelA-modulating nanobodies Nb94 and Nb96 could have an impact on the virulence of *E. coli* mimicking the effects of genetic knockouts catalytically impaired variants of RelA.

We used the wax moth *Galleria mellonella* larvae infection model to assess this effect on the *E. coli* clinical isolate Ec156^45^. In the presence of the control Nb120 (produced by the *Ec156_nb120* strain), which was not selected against RelA, only 30% of the larvae survived the infection after 36h. By contrast when the *Galleria* larvae were challenged with the *Ec156_nb94* and *Ec156_nb96* (Ec156 strains^45^ transformed with plasmids expressing the Nb94 and Nb96 nanobodies) survival increased to 50% and 70% respectively (**Fig. 1e**). Notably, these strains displayed no growth defects when grown on M63 plates.

Our infection assays demonstrate that while *E. coli* can tolerate the loss of activity of RelA when grown in nonstressed laboratory conditions without a dramatic fitness defect, a fully functional (p)ppGpp synthetase is crucial for a successful infection.

### Nb94 and Nb96 block the TGS of RelA_*Ec*_

To gain a functional insight into the nanobody’s mode of action, we have used AlphaFold^46^ to model the structure of the RelA*_Ec_*:Nb94 and RelA*_Ec_*:Nb96 complexes. High confidence models predict that Nb94 and Nb96 recognize RelA*_Ec_ via* the TGS through a ≈990 Å^2^ and ≈850 Å^2^ interface respectively (**Fig. 1f-g**). The complementarity-determining regions 2 and 3 of Nb94 and Nb96 provide the majority of the contacts (with CDR1 not involved in binding) by interfacing with the Threonyl-tRNA Synthetase, GTPase and SpoT (TGS) domain of RelA through the central α-helical region and the C-terminal β-strand, as well as via additional contacts with the α6-α7 motif of the pseudo-HD domain and the Core domain. Interestingly, the complex interface is dominated by the H432 residue located in an antiparallel α-helical motif of the TGS that makes extensive contacts with CDR2 and CDR3 and β-strands β3 and β4 Nb94 (**Supplementary Fig. 1a**).

In the case of Nb96, the AlphaFold-predicted mode of TGS recognition differs. Nb96 engages the opposite face of the antiparallel α-helical motif within the TGS domain through all three CDRs, and additionally establishes extensive contacts via β-strands β3, β4, and β5. Notably, both nanobodies interact directly with residue H432, although in the case of Nb96, this interaction occurs specifically through CDR2. (**Fig. 1g** and **Supplementary Fig. 1b**). These differences in recognition reflect substitutions within the complementarity-determining regions, particularly in CDR2 (**Supplementary Fig. 1c**). Nevertheless, as both Nb94 and Nb96 nanobodies target the TGS domain of RelA, they are likely to disrupt the allosteric crosstalk between the catalytic and regulatory regions. Furthermore, their binding provides a structural basis for the potent inhibition of synthesis observed with Nb94 and Nb96, as they are predicted to tether the TGS domain to the pseudo-HD and Core domains, thereby likely preventing the enzyme from adopting its active, open conformation^26,27^, while also blocking the recognition of uncharged tRNAs on the A site by the TGS domain which is mediated by H432^47^. We used ITC to monitor the interaction of Nb94 and Nb96 with an H432E-substituted version of RelA*_Ec_* (RelA*_Ec_*^H432E^), a well characterized variant that abrogates RelA’s functionality by abolishing RelA activation by deacylated tRNA^17,47^. In the case of the interaction of RelA*_Ec_*^H432E^ with Nb94 and Nb96, the introduction of a negative charge to the interface is predicted to disrupt binding. Indeed, the lack of binding of either nanobodies to RelA ^H432E^ (**Supplementary Fig. 1d-e**) supports the proposed mode of recognition and inhibition of ppGpp synthesis through sequestration of the TGS domain.

### Nanobodies modulate Rel_*Ct*_ catalytic activities

While the monofunctional SYNTH-only RelA can only explore the resting and extended states observed in long RSH ppGpp synthetases, bifunctional SYNTH- and HD-competent Rel enzymes have a much broader conformational landscape^18^. Being capable of both, hydrolysis and synthesis of (p)ppGpp, these proteins readily explore different conformational states^18^. These states involve a compact HD^ON^-compatible τ-state, in which the NTD are close together to with synthetase domain completely occluded. The enzyme also explores a relaxed state (less structured in comparison with the τ-state) in which both catalytic domains are partially open and marginally active; and an elongated, ribosome-bound state, SYNTH^ON^-compatible, in which the NTD is fully open and the hydrolase active site is disordered and inactive^18,19^. Therefore, discovering and engineering specific conformational modulators of Rel is in principle more complex. We turned to the bifunctional (p)ppGpp synthetase/hydrolase Rel from the model thermophile *C. tepidum* (Rel*_Ct_*) to test whether nanobodies could selectively restrict the allosteric landscape of long RSH factors to trap the enzyme in a specific state that would define its catalytic output.

Using the same strategy to identify nanobodies targeting RelA*_Ec_*, we selected two families of nanobodies – Nb898 and Nb585 – that strongly bind Rel*_Ct_* (**Supplementary Fig. 2**). ITC showed that full-length Rel*_Ct_* interacts with Nb898 and Nb585 with *K*_d_ values of 186 nM and 21 nM, respectively (**Fig. 2a-b** and **Supplementary Table S1**). In both cases this interaction is confined to the N-terminal catalytic region (NTD) of the enzyme since C-terminal truncations did not affect binding (**Supplementary Table S1**). In the absence of Rel SYNTH stimulators, pppGpp or 70S *E. coli* ribosomes, Rel*_Ct_* is a poor catalyst of ppGpp synthesis (**Fig. 2c**). Addition of *E. coli* 70S ribosomes and pppGpp increases the SYNTH activity ≈8 times. In the presence of Nb898, the turnover of Rel*_Ct_* increased ≈11-fold, on par with the enhancing effects of the 70S and pppGpp combined. This effect is accompanied by a 1.6-fold decrease in hydrolase activity (**Fig. 2d**) that results in the accumulation of ppGpp. By contrast, Nb585 turns SYNTH^OFF^, strongly inhibiting (p)ppGpp synthesis (**Fig. 2c**) while increasing the hydrolase activity of Rel*_Ct_* by two-fold (**Fig. 2d**). These results are consistent with the nanobodies modulating the antagonistic allosteric coupling between the two catalytic domains of Rel*_Ct_* to control the enzymatic output.

**Fig. 2.**
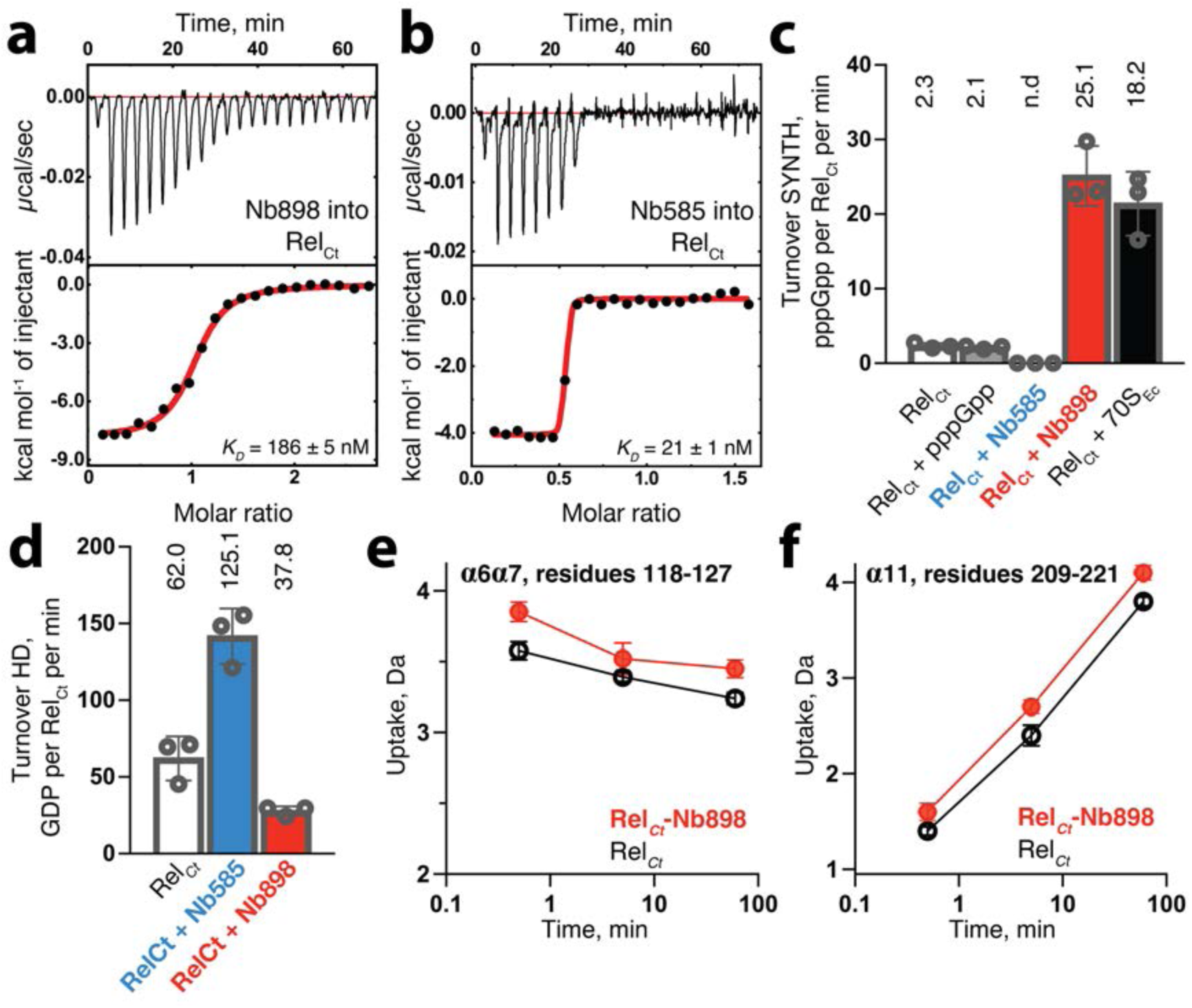
Nanobodies targeting Rel can modulate the activity of the enzyme by restricting their allosteric landscape. ITC titrations of Nb898 **(a)**, Nb585 **(b)**, into Rel*_Ct_*. **(c)** SYNTH activity of Rel*_Ct_* unbound (empty bar); in the presence of Nb898 (red bars); Nb585 (blue bars); pppGpp (grey) and 70S ribosomes (black). **(d)** HD activity of Rel*_Ct_* unbound, in the presence of Nb898 (red bars), Nb585 (blue bars). The contrasting features of the action of the nanobodies in (c) and (d) suggest they highjack the enzyme’s conformational landscape to favour one activity in detriment of the other. **(e)** Deuterium uptake profiles for representative peptides the allosteric hotspot region involving residues 118-127 of α6α7 **(e)** and residues 209-221 of α11 **(f)**.

### Nb898 increases Rel ^NTD^ dynamics

We monitored the structural effects of Nb898 binding using Hydrogen Deuterium eXchange coupled with Mass Spectrometry (HDX-MS) by comparing Rel ^NTD^ (a more stable and soluble fragment that contains the Nb binding site) in its substrate-free unliganded state, and in the Rel ^NTD^:Nb898 complex (**Supplementary Fig. 3a**).

The overall effect of Nb898 binding was an increased deuterium uptake (ΔHDX) involving both catalytic regions that radiates from the central region of the enzyme (**Fig. 2e-f** and **Supplementary Fig. 3a**). The highest uptake increase localizes to the allosteric hub involving the α6α7 active site motif (residues 101 - 135) and α11 (residues 201 - 236) that mediate the coupling between SYNTH and HD domains^17,19^. The higher degree of uptake observed in this crucial allosteric region (**Fig. 2e-f**) suggests that the binding of Nb898 triggers a more dynamic state of the enzyme in which the SYNTH active site is more exposed and primed for substrate binding and catalysis. This aligns closely with prior structural evidence showing that α11 unwinds in Rel and RelA upon activation of the SYNTH domain^17,19^.

The HDX-MS data also detected a prominent decrease in deuterium exchange in the region that contains α-helix α13 (residues E279 to Y290), which is indicative of the direct association of Nb898 with Rel ^NTD^ and showed a slight additional deuterium protection signal in the hydrolase active site (residues 65-78, **Supplementary Fig. 3a**). A more moderate protection was observed towards the C-terminal in the region. This involved Rel’s G-loop (residues 304 - 316, **Supplementary Fig. 3a**), suggesting a possible allosteric stabilization of the residues involved in GDP/GTP binding that could stimulate catalysis.

To confirm the binding interface, we probed the epitope define by α13 using point substitutions in Rel*_Ct_* (K280A, D283A, F285A, A286K, Y290A) to monitor the interaction with Nb898 by ITC. In all cases we detected an increase in the binding *K*_d_ of the Rel*_Ct_* variants compared to the WT Rel*_Ct_* confirming the binding epitope (**Fig. 3a-d** and **Supplementary Table S1**). Collectively these results indicated that the direct interaction of Nb898 with the SYNTH domain triggered the closing or rigidification of HD and consequent inactivation of the hydrolase function and activated the SYNTH function by allosterically increasing backbone dynamics.

**Fig. 3.**
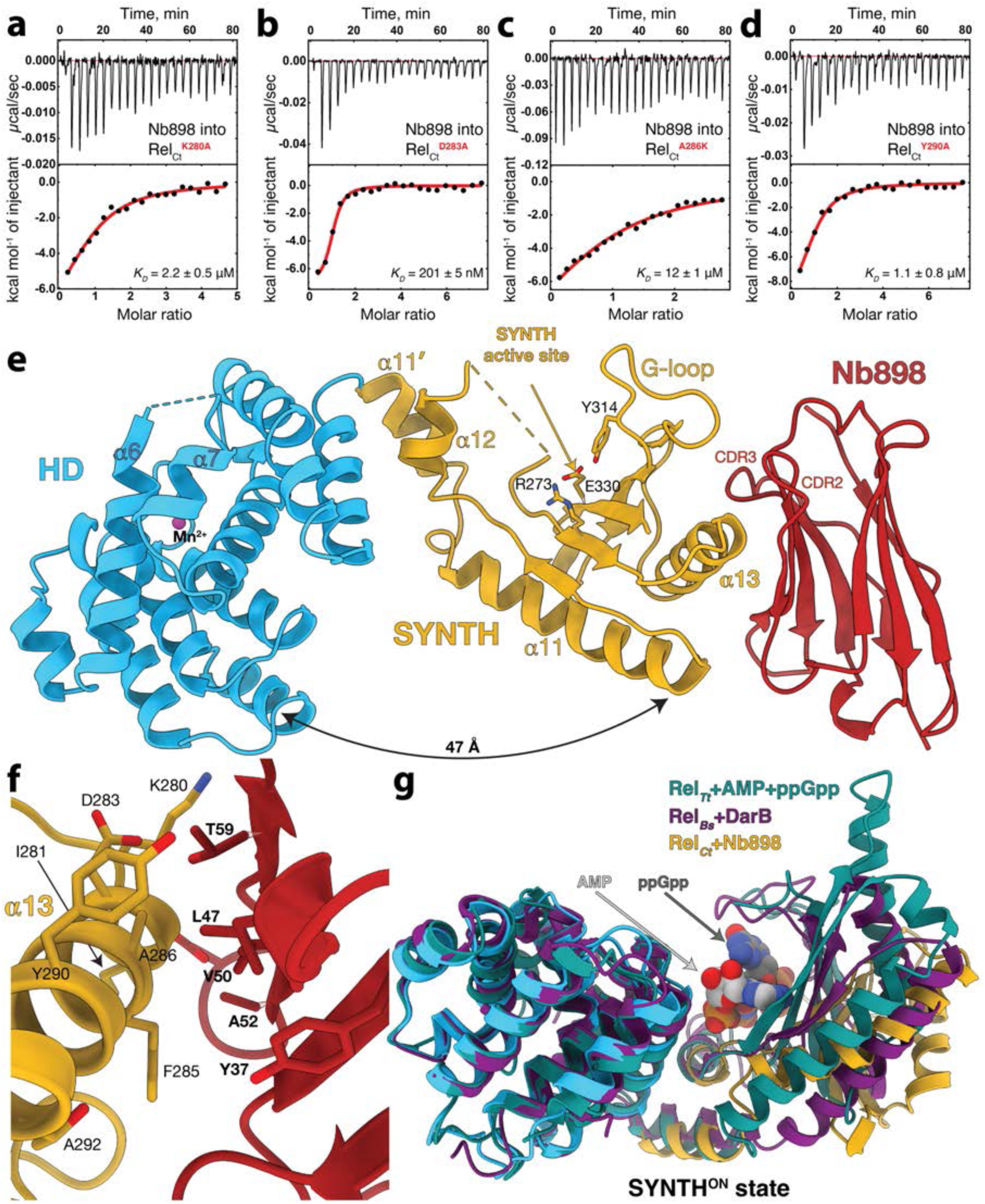
Nb898 binds to the SYNTH domain of Rel*_Ct_* contacting α13 and the G-loop. ITC titrations of Nb898 into Rel*_Ct_*^K280A^ **(a)**, Rel*_Ct_*^D280A^ **(b)**, Rel*_Ct_*^A286K^ **(c)**, and Rel*_Ct_*^Y290A^ **(d)**. into Rel*_Ct_* validate the binding interface. **(e)** Crystal structure of the Rel*_Ct_*^NTD^-Nb898 complex. Rel’s HD and SYNTH domains are coloured as in Fig. 1f, and Nb898 in red. CDR2 and CDR3 are labeled. **(f)** Details of Rel*_Ct_*^NTD^-Nb898 binding interface highlighting key structural elements. **(g)** Superposition of the catalytic domains of Rel*_Ct_* onto the Rel*_Tt_*-AMP-ppGpp complex (PDBID 6S2U, in green) and the Rel*_Bs_*-DarB complex (PDBID 8ACU, in purple). The superposition confirms Nb898 selects the open, SYNTH^ON^ conformation of the enzyme. The AMP and ppGpp nucleotides observed in the post-catalytic state of Rel*_Tt_* are are highlighted with light and dark grey arrows respectively.

### Nb898 stabilizes the SYNTH^ON^ active state of Rel_*Ct*_^NTD^

In order to understand the effect of Nb898 on catalysis, we determined the structure of the Nb898:Rel*_Ct_*^NTD^ complex (**Fig. 3e**). Inspection of the complex confirms the binding interface predicted by the HDX-MS: indeed, Nb898 engages Rel*_Ct_*^NTD^ via the SYNTH domain. The interactions from the nanobody side occur mainly through the core β-sheet of the Ig-domain, CDR2 which forms a flat hairpin structure with a pronounced bulge in the middle stacking α13, plus additional contribution from CDR3 (**Fig. 3e**). This binding interface is in perfect agreement with the HDX and mutagenesis results (**Fig. 2e** and **3a-d**). The stacking interface is largely hydrophobic, composed by the side chains of residues I281, F285, A286, Y290 and T292 from the Rel*_Ct_* side and Y37, A47, V50, A52, D55 and T59 from Nb898. In addition, T59 forms a strong hydrogen bond with D283 tethering CDR2 to α13 and the backbone of T58 hydrogen-bonds K280 (**Fig. 3f**).

In the conformation of Rel*_Ct_*^NTD^ induced by Nb898, the HD and SYNTH domains are distanced 47 Å (**Fig. 3e**). The active site of the HD domain is largely misaligned with the ED catalytic motif away from the Mn^2+^ ion and the pocket that coordinates the 3’-pyrophosphate of (p)ppGpp during hydrolysis. By contrast, the SYNTH domain active site and catalytic residues are exposed to the solvent. This is accompanied with the fracturing of α11 (into α11’ and α11) and the displacement of α12, all signature structural elements of the activation of catalysis by the SYNTH domain (**Fig. 3e**). Noteworthy, this mode of recognition via α13 mimics that of *Thermus thermophilus* Rel (Rel*_Tt_*) bound to the reaction products ppGpp and AMP^19^ (**Fig. 3g**) as well as of *B. subtilis* Rel (Rel*_Bs_*) bound DarB in absence of cAMP^48^ (**Fig. 3g** and **Supplementary Fig. 3b-c**); the latter also mediates the activation of ppGpp synthesis and inhibition of hydrolysis^48^ (**Fig. 3g**).

Catalysis by long RSH enzymes is directly linked to their conformational state^18,27^. Binding of GDP and ATP trigger the opening of the NTD region exposing the SYNTH active site while binding of ppGpp exposes the HD domain and inactivates SYNTH^19^. Using GDP and APCPP to induce the active SYNTH state of Rel*_Ct_*^NTD^, we determined the structure of the substrate-liganded complex at 2.65 Å. The structure shows the binding of both nucleotides, indeed, promotes the open conformation (**Fig. 4a** and **Supplementary Fig. 3d-e**). The guanosine base of GDP is stacked by the G-loop Y314 and the 5’-diphosphates stabilized by residues K255, K261, and K303. APCPP is held deep into the active site (**Fig. 4b**). The adenine base is strongly coordinated by R245 and R273 whereas the triphosphate moiety is locked in position for catalysis by R245, K247, S251 and K255, bringing the β-phosphate close to the 3’-OH of GDP (**Fig. 4b**). This open state bound to the substrates matches strongly the conformation of Rel*_Ct_*^NTD^ observed in the complex with Nb898 (**Fig. 4a**). Collectively these results indicate that Nb898 triggers the SYNTH^ON^ conformation of Rel ^NTD^ even in absence of substrates.

**Fig. 4.**
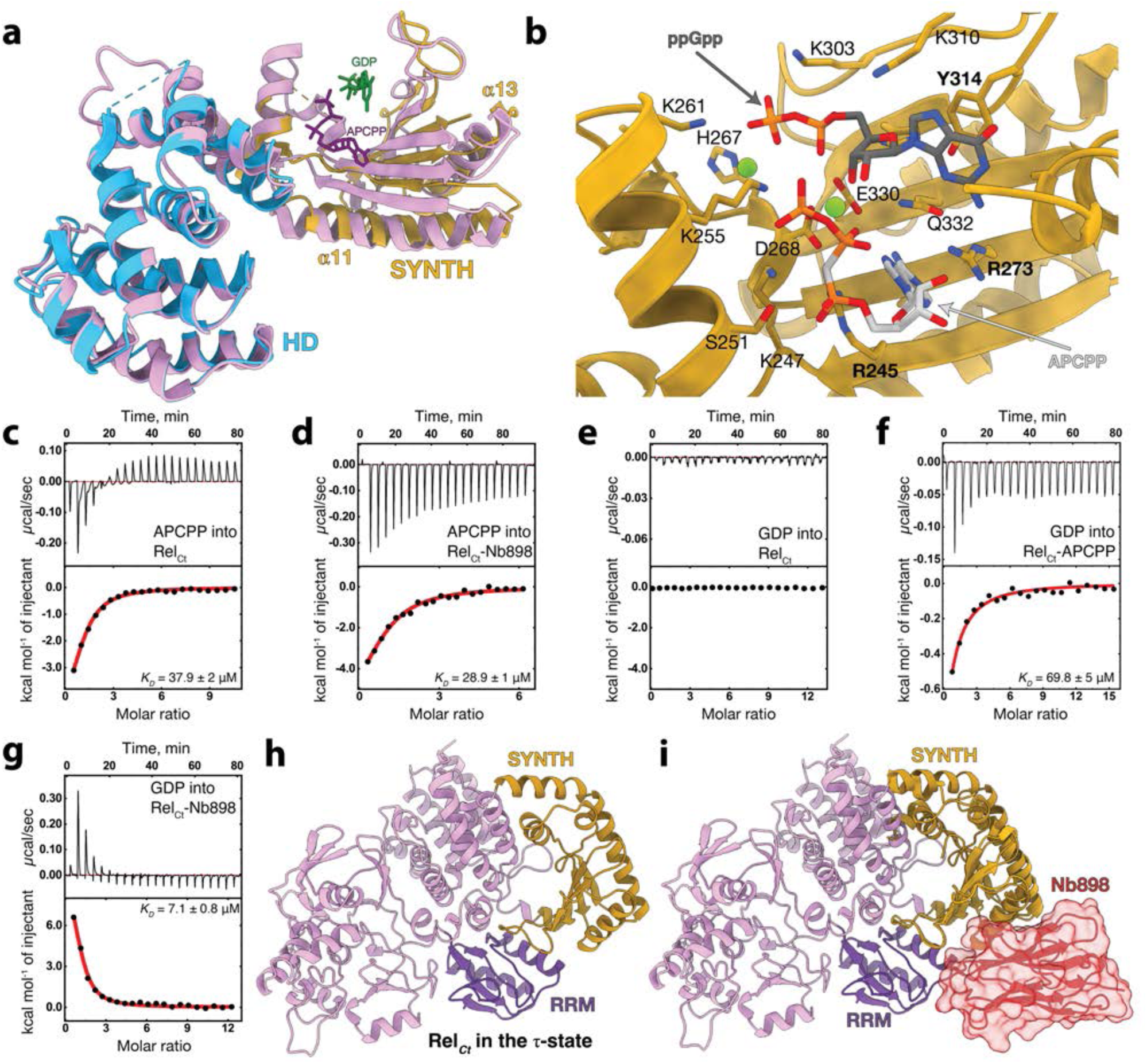
Nb898 binds to the SYNTH domain of Rel*_Ct_* and allosterically enhances the affinity of the enzyme for GDP. **(a)** Superposition of the structure of the Rel*_Ct_*^NTD^-Nb898 complex (coloured as in Fig. 1f) on the structure of Rel*_Ct_*^NTD^ (in pink) in complex with APCPP (in purple) and GDP (in green). Crucial structural elements involved in the ON/OFF switch of the enzyme such as α11, α13 are labeled in the figure. The superposition confirms Nb898 triggers an active state of Rel*_Ct_*^NTD^ compatible with nucleotides binding. **(b)** Details of Rel*_Ct_*^NTD^-APCPP-GDP complex interface highlighting key active site residues involved in substrate binding and catalysis. **(c)** ITC titration of APCPP into Rel*_Ct_*^NTD^; **(d)** APCPP into Rel*_Ct_*^NTD^-Nb898 complex; **(e)** GDP into Rel*_Ct_*^NTD^; **(f)** GDP into Rel*_Ct_*^NTD^-APCPP complex; **(g)** GDP into Rel*_Ct_*^NTD^-Nb898 complex. **(h)** Cartoon representation of a model of full-length Rel*_Ct_* in the hydrolase-compatible τ state. The SYNTH (in gold) and RRM (purple) domains are labeled in the figure. **(i)** Surface representation of an idealized (unrealistic) model of full-length Rel*_Ct_*^NTD^ in complex with Nb898 (in red). The model confirms that the Rel*_Ct_*^NTD^-Nb898 complex is incompatible with the hydrolase-active τ state.

### Nb898 stimulates SYNTH activity Rel ^NTD^ by promoting nucleotide binding

RSH enzymes typically display a preferential order for the incorporation of substrates to the SYNTH active site^19,48,49^. We used ITC to probe the incorporation of nucleotides to Rel ^NTD^. Rel ^NTD^ binds the APCPP nonhydrolyzable ATP analogue with an affinity of 37.9 μM, which is enhanced by 1.3-fold in the presence of Nb898 (**Fig. 4c-d**). By contrast, in the absence of APCPP or Nb898, we were not able to detect binding of GDP to Rel ^NTD^ (**Fig. 4e**). In the presence of saturating APCPP concentration (5-fold above the *K*_d_) the enzyme binds the GDP substrate with a *K*_d_ of 69.8 μM, and Nb898 has a 10-fold stimulatory effect on the interaction (*K*_d_ = 7.0 μM) (**Fig. 4f-g**). These results indicate that Rel ^NTD^ binds its substrates in a sequential and ordered manner, initiated by ATP and followed by GDP or GTP. In addition, they suggest that the stabilizing presence of Nb898 enhances the affinity of the enzyme for both nucleotides, which likely underlines the SYNTH-stimulatory effect of the nanobody.

### Nb898 modulates Rel_*Ct*_ τ-state precluding CTD autoinhibition and HD activation

To further investigate the mechanism of allosteric activation of Rel*_Ct_* by Nb898, we modeled the structure of unliganded full-length Rel*_Ct_* based on the structure of the highly compact τ-state of *A. baumannii* SpoT (SpoT*_Ab_*) which represents a CTD-autoinhibited long RSH primed for hydrolysis (**Fig. 4h**). Indeed, the integrity of the association of the CTD of long RSHs with its HD and SYNTH domains is crucial for the stability of the SYNTH-inhibited τ-state (SYNTH^OFF^) of SpoT and Rel as evidenced by a decrease in their hydrolase activity upon destabilizing mutations or domain deletions^18,25,50^. This contrast with the SYNTH activity: the mechanism of activation of Rel, and the role of the CTD in preventing this activity, are only relevant in the context of starved ribosomes with the CTD autoinhibition being only a part of a process that also involves the activation and stabilization of the fully active enzyme by A-site tRNAs and pppGpp. While the CTD itself is intricately involved in ribosome binding, in the absence of ribosomes, CTD truncations do exhibit an increase in ppGpp synthesis^20,25^. In addition, as observed for the Rel activator DarB^48^, a ribosome-independent stabilization of an open, elongated state of Rel could have an impact in the SYNTH activity of the enzyme^48^.

When we compared the structure of the Rel*_Ct_*-Nb898 complex with the model of SYNTH^OFF^ Rel*_Ct_* it becomes apparent that both states are incompatible. The strong binding of Nb898 would clearly interfere with the RRM:SYNTH association which would drive the conformational equilibrium of the enzyme toward a relaxed non-autoinhibited state more compatible with SYNTH activity (**Fig. 4i**). Collectively these results suggest that Nb898 hijacked a naturally existing allosteric pathway of activation of long RSH factors. In this way, Nb898 restricted the conformational landscape of the enzyme to the SYNTH open state, allosterically priming catalytic residues and enhancing substrate recognition. This resulted in an increased ppGpp synthesis.

### Nb585 locks Rel_*Ct*_ and primes the HD domain while switching off the SYNTH domain

As with the SYNTH-activating Nb898, we used HDX-MS to characterize the interaction of the HD-activating Nb585 with Rel*_Ct_*. In the presence of Nb585, we observed deuterium protection signal clustering in the central region of the enzyme that comprises the C-terminal part of the HD domain that mediates the coupling between SYNTH and HD domains and α11. These structural elements are all solvent exposed in the open pre-catalytic state of Rel*_Ct_* when bound to GDP and APCPP and in the complex with Nb898. However, these regions coalesce together around the central axis of the enzyme in a closed state and become protected from deuterium exchange (**Fig. 5a-b**). By contrast in the HD domain, the active site shows moderate increase in deuterium uptake consistent. This becomes apparent in the regions containing K47-R48 involved in guanosine coordination and H57, H82, D83 involved in Mn^2+^ coordination (**Fig. 5a and c-d**). Collectively these results are consistent with the exposure of catalytic residues upon binding to Nb585 and the stabilization of a HD^ON^ state.

**Fig. 5.**
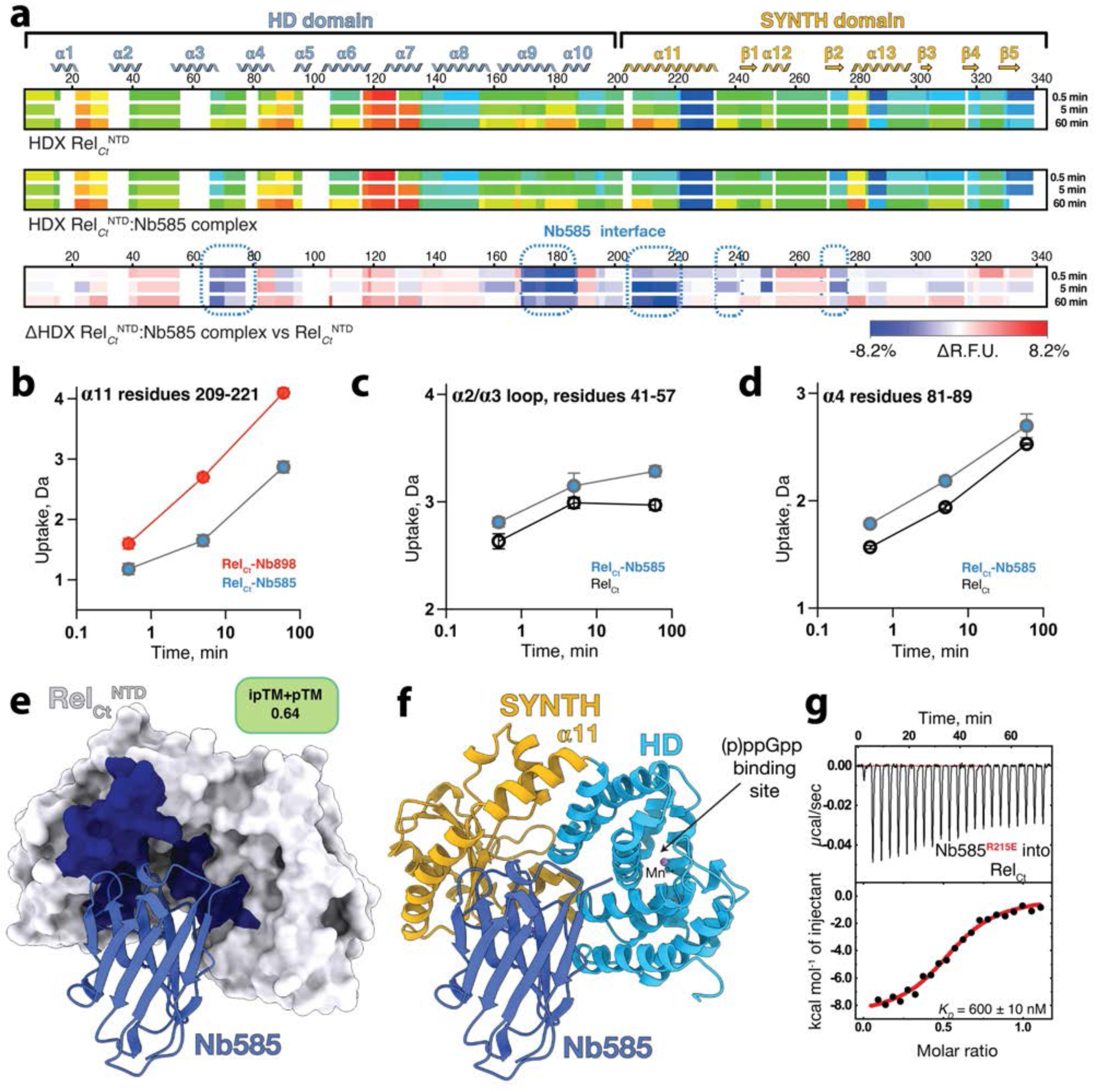
Nb585 contacts both HD and SYNTH domains of Rel*_Ct_* stabilising a closed conformation that is compatible with ppGpp hydrolysis. **(a)** Heatmaps representing the HDX of Rel*Ct*^NTD^ (top), Rel*Ct*^NTD^-Nb585 complex (center) and ΔHDX (bottom). Residues involved in the binding interface are outlined by a dashed light blue line. **(b)** Deuterium uptake profile for a representative peptide the allosteric hotspot region involving residues 209-221 of α11 of Rel*Ct*^NTD^-Nb585 complex in blue vs Rel*Ct*^NTD^-Nb898 in red. Deuterium uptake profile for a representative peptide of the α6/α7 loop **(c)** and the active site fragment of α4 **(d)**. **(e)** Surface representation of Rel*Ct*^NTD^ (in light grey) bound to Nb585 (in blue) as predicted by AlphaFold. Rel*Ct*^NTD^ residues involved in the binding interface are coloured in dark blue. **(f)** Cartoon representation of the model of Rel*Ct*^NTD^-Nb585 complex with Rel*Ct*^NTD^ HD and SYNTH domains coloured as in Fig. 1f and Nb585 in blue. The Mn2+ ion, a proxy for the (p)ppGpp binding site in the HD domain is highlighted in the figure. **(g)** ITC titration of Nb585 R215E substituted version (Nb585^R215E^) into Rel*Ct*^NTD^. This position is predicted by AlphaFold and HDX-MS to be involved in the complex interface. The increase in K*D* triggered by the R215E substitution supports the predicted mode of action of Nb585. RFU, relative fractional uptake.

The HDX-MS data are in good agreement with the AlphaFold prediction of the structure of the complex that places Nb585 between HD and SYNTH domains. In the predicted complex Nb585 interacts with the SYNTH domain via the region involved in ATP, reminiscent of the way toxic Small Alarmone Synthetases (toxSAS) are inhibited by their antitoxins^51^, which is consistent with the strong inhibition of synthesis displayed by Nb585. This binding mode tightly tethers the HD and SYNTH domains (**Fig. 5e-f**) stabilizing a closed state that resembles that of Rel*_Tt_* bound to ppGpp^19^ (*r.m.s.d* of 0.9 Å, **Supplementary Fig. 4a**). According to the model and consistent with the HDX-MS data, Nb585 makes extensive contacts with the C-terminal α-helix of the HD domain and α-helix α11 of SYNTH. In particular, R215 of Rel*_Ct_* is predicted to form a salt bridge with D31 of Nb585 (**Supplementary Fig. 4b-c**). This region contains the kink implicated in the closed/open switch of the NTD and is strongly protected from deuterium exchange upon Nb585 binding (**Fig. 5a-b**). To challenge this, we introduced the R215E substitution in Rel*_Ct_* and observed a 28-fold drop in affinity of Nb585 for Rel*_Ct_*^R215E^ (**Fig. 5g**). This is consistent with our model of the complex (**Fig. 5e**) and the HDX-MS and biochemical data (**Fig. 2c-d** and **5a**). Collectively these results suggest that long RSH-binding molecules that activate one catalytic function likely trap the enzyme in a restricted conformational state that precludes the opposing catalysis.

## Discussion

The stringent response is a key regulatory pathway involved in virulence and antibiotic resistance. Thus, targeting the stringent response via chemical inhibition of RSH enzymes is an attractive strategy for development of antibacterials that disrupt the (p)ppGpp homeostasis. Here we identified nanobodies that are efficient allosteric modulators of long SYNTH-competent ribosome-associated long RSH Rel and RelA and dissected their mechanism of action. We showed that these nanobodies are effective tools to control the catalytic output of these enzymes *in vitro* and *in vivo* precluding bacterial growth under nutrient starvation.

Long RSH enzymes are naturally controlled by adaptor proteins that modulate the cellular levels of (p)ppGpp alarmones in addition to the canonical ribosome-dependent pathway (**Fig. 6a-c**). In *E. coli*, Rsd is a stimulator of the (p)ppGpp-hydrolase activity of SpoT during carbon source starvation^32^ (**Fig. 6a**). In absence of c-di-AMP, DarB binds strongly to the SYNTH domain of *B. subtilis* Rel^48^ while analogously, upon phosphorylation, *C. crescentus* EIIA^Ntr^ binds the RRM domain of Rel which leads to a strong inhibition of hydrolysis^31^ and ppGpp accumulation during nitrogen starvation^30^. Both regulators preclude the formation of the τ-state of Rel^18^ and inhibit the hydrolase function by restricting the conformational space to an open, SYNTH^ON^ conformation only compatible with (p)ppGpp synthesis^31,48^. The combined allosteric effect of favoring synthesis while strongly inhibiting hydrolysis leads to the accumulation of (p)ppGpp. By contrast, NirD binds to both domains of RelA^NTD^ and prevents (p)ppGpp synthesis^36^ (**Fig. 6b-c**).

**Fig. 6.**
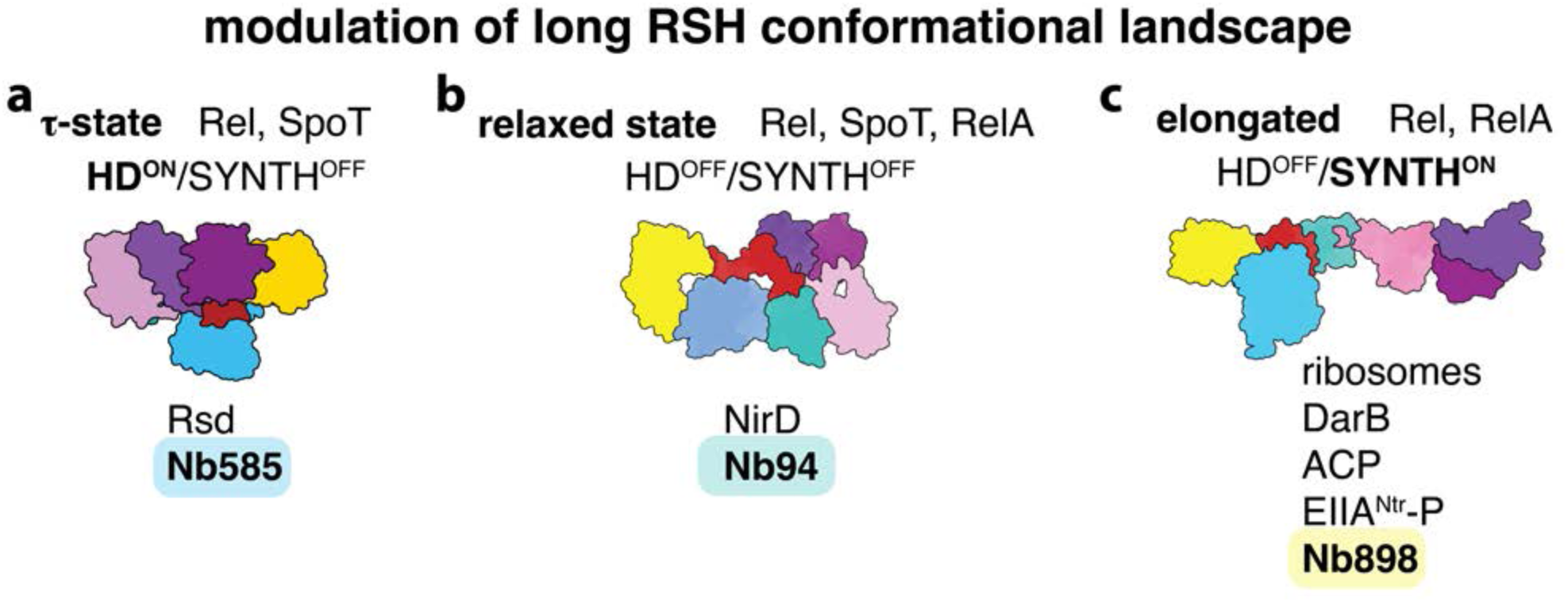
Naturally occurring allosteric sites of RSH enzyme can be exploited to modulate their activity. Nanobodies such as Nb585 **(a)**; Nb94 **(b)**; and Nb898 **(c)**, bind to Rel and RelA mimicking the mode of action of endogenous RSH adaptor proteins that are activated during the cell cycles to modulate the metabolic state. The action of these nanobodies suggests that besides the genetic well-characterised genetic effects on antibiotic tolerance, virulence and survival; chemically targeting RSH enzymes could be a viable pathway to develop drugs active against bacteria.

We demonstrated that the nanobodies characterized in this study target key regulatory hotspots in long RSH enzymes, acting as catalytic checkpoints. Nb94 and Nb96 recognize and occlude the antiparallel α-helical motif of the *E. coli* RelA TGS domain – an element that inspects the CCA end of tRNAs and is essential for RelA’s activation on the starved ribosomal complex^26,27^. Thus, selective inactivation the TGS domain with Nb94 and Nb96 countered the intramolecular allosteric signaling that triggers RelA-mediated (p)ppGpp synthesis in response to amino acid starvation.

Analogously, Nb898 binds the structural epitope that we have earlier shown in the case of *B. subtilis* Rel to be recognized by SYNTH-stimulatory factor DarB^48^. Neither Nb898 nor DarB make direct contacts with the active site of the enzyme to enhance (p)ppGpp synthesis. Rather, both modulators counter the formation of the hydrolase-compatible τ-state by preventing the interaction between the RRM domain and the SYNTH domain. This, in turn, leads to a partial opening of SYNTH (**Fig. 4h-i**). Indeed, binding of Nb898 enhances the affinity of the enzyme for ATP, mimicking the stimulatory effect of the 70S ribosome and resembling the genetic truncation of the RRM domain of Rel and RelA that induces slow growth by the toxic accumulation of (p)ppGpp^20^ (**Fig. 6c**). The stronger stimulatory effect of Nb898 compared to DarB is likely the result of the higher affinity of Nb898 for Rel (186 nM in the case of Nb898 *vs* 1.4 μM in the case of DarB^48^), suggesting that tight-binding molecules targeting this particular region of Rel are likely to induce severe growth arrest. By contrast, Nb585 recognizes a more complex tridimensional epitope that includes both N-terminal catalytic domains. This is reminiscent of the conformational recognition employed by NirD^36^ to inhibit RelA. However, from a functional perspective, Nb585 mimics Rsd^32^ by stabilizing the close τ-state of the long RSH enzyme to increase the hydrolase activity while inhibiting the (p)ppGpp synthesis.

Collectively, functional effects mediated by nanobodies mirror the activity-defining rearrangements stabilized by nanobodies on the cystic fibrosis transmembrane conductance regulator (CFTR)^39,52^ and GPCRs^53^. Nanobodies targeting extracellular epitopes of proteins are currently being developed as potential drugs for a variety of human diseases, including antimicrobials, however, they are mainly targeting secreted toxins and toxin receptors. While practical applications of intracellular delivery translated to therapeutics are likely to remain a significant challenge, the strong effect shown by the action of the nanobodies on the *Galleria* infection model suggest the RSH pathway is a viable target to develop novel antibacterials. Rather, small molecule compounds are still the method of choice for intracellular therapeutic targets. Thus, the tridimensional allosteric hotspots revealed using these nanobodies, provides valuable structural insights that can guide drug development and inform future studies aimed at modulating the enzymatic output of long RSHs.

## Acknowledgments

This work was financially supported by the Fonds National de Recherche Scientifique (CDR J.0065.23F; PDR T.0090.22), ERC (CoG DiStRes, n° 864311), the Fonds Jean Brachet, the Fondation Van Buuren and FNRS WELBIO ADV X.1520.24F, INSTRUCT (PID: 4098), all to A.G-P. C.M. was supported by Fonds de la Recherche Scientifique (MIS grant F.45322.22). K.C.W. and K.V.N. were supported by FRIA. The authors acknowledge the use of the PROXIMA-2A beamline at the Soleil synchrotron (Gif-sur-Yvette, France) and I24 (Diamond Light Source synchrotron Oxfordshire UK). V.H. was supported by the Knut and Alice Wallenberg Foundation project grant 2020-0037 to V.H., Vetenskapsrådet grants 2021-01146 and ÄR-MH 2024-06059 as well as Göran Gustafsson Foundation for Research in Natural Sciences and Medicine (the Göran Gustafsson Prize to V.H.).

## Supplementary Figures

**SFig. 1.**
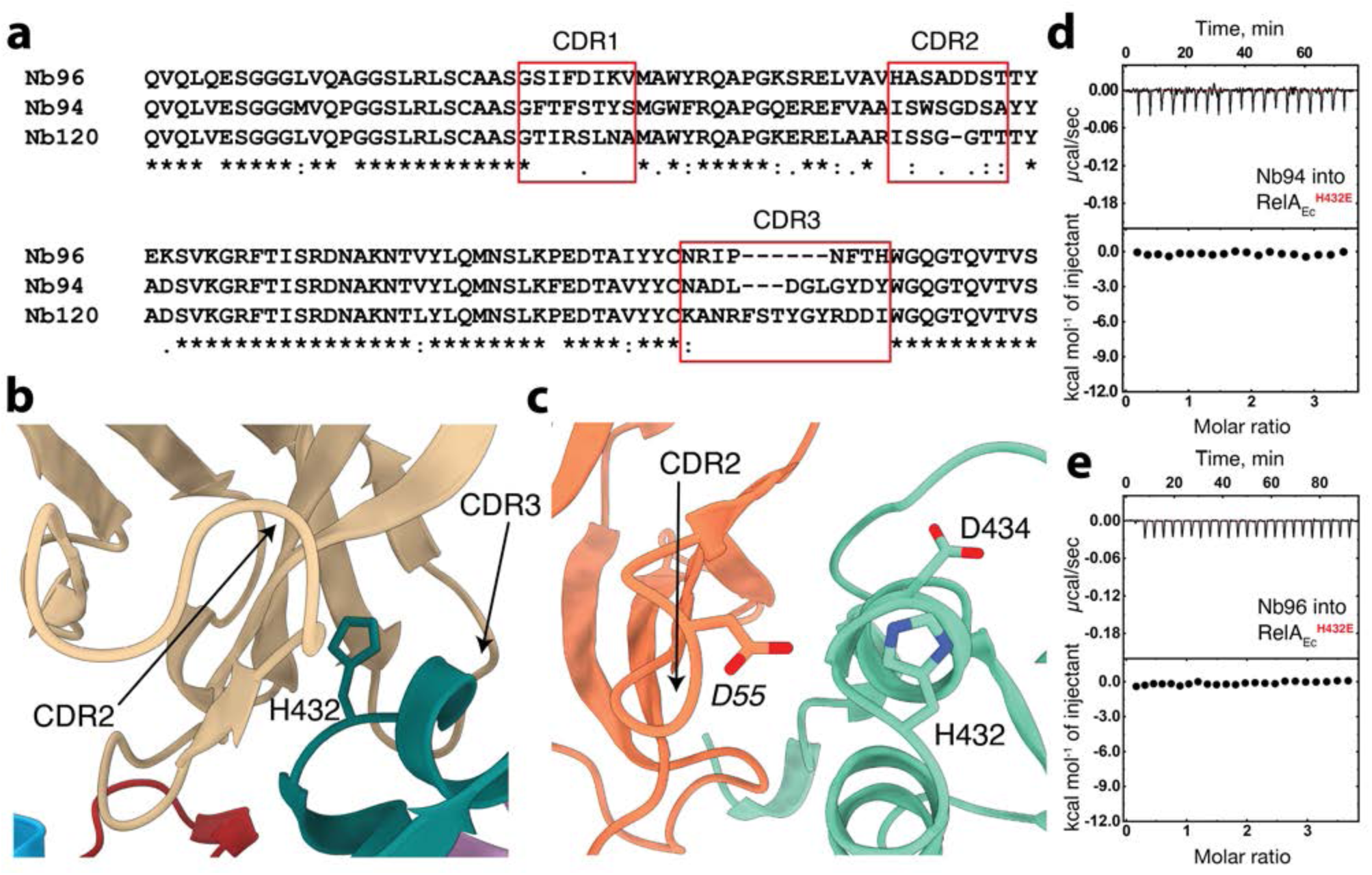
The TGS H432 directly coordinates Nb94 and Nb96. **(a)** Multiple sequence alignment of the nanobodies Nb94, Nb96 and the control Nb120 (each CDR region is boxed and labeled). Details of the binding interface of Nb94 **(b)** and Nb96 **(c)** as predicted by AlphaFold. CDR regions involved in binding are indicated by a black arrow. From the prediction it is clear the H432E substitution will significantly disturb the interface. ITC titrations of Nb94 **(d)** and Nb96 **(e)** into RelA*_Ec_*^H432E^. The lack of binding observed strongly supports the predicted mode of binding.

**SFig. 2.**
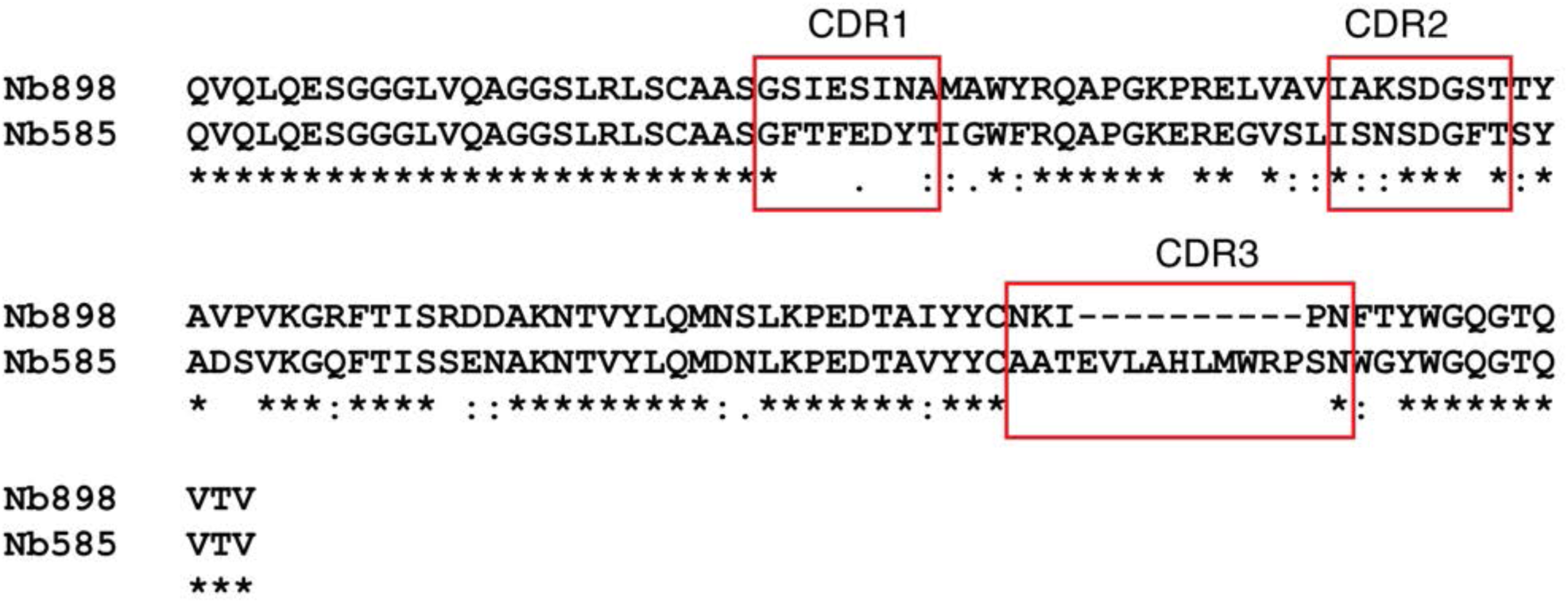
Nb898 and Nb585 have notably different CDR regions. Sequence comparison of Nb898 and Nb585 shows the strong differences in the observed complementarity determining region (CDR) that defined the contrasting activities of these nanobodies. Each CDR region is labeled in the figure.

**SFig. 3.**
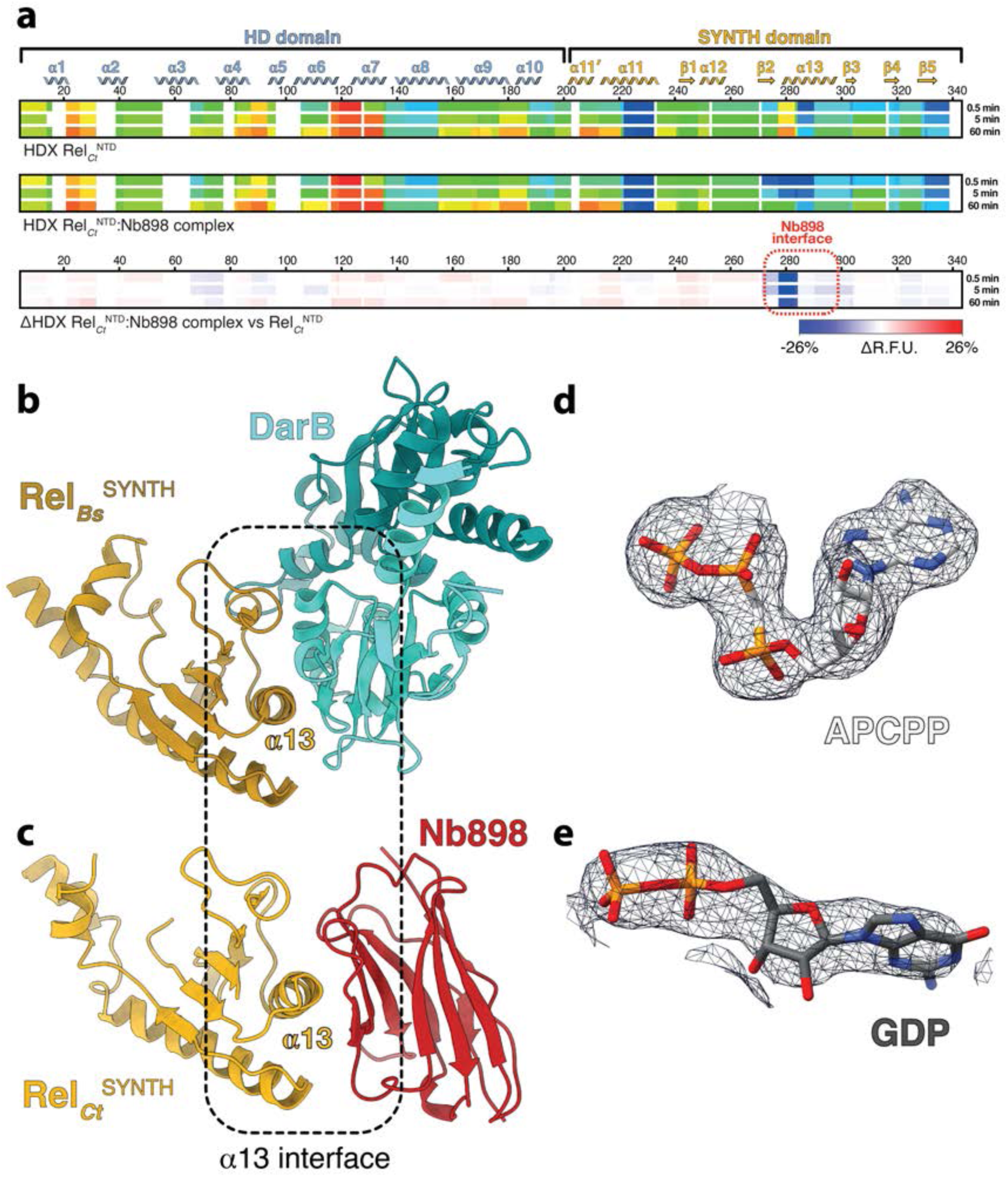
Characterisation of the Rel_Ct_^NTD^-Nb898 interface. **(a)** Heatmaps representing the HDX of Rel_Ct_^NTD^ (top), Rel_Ct_^NTD^-Nb898 complex (center) and ΔHDX (bottom). Residues involved in the binding interface are outlined by a dashed red line. RFU, relative fractional uptake. DarB **(b)** and Nb898 **(c)** enhance the (p)ppGpp synthetase activity of Rel. In both cases α13 is the main contact point with additional interactions contributed by α11 and the active site G-loop. Unbiased mFo-DFc electron density map corresponding to the bound APCPP **(d)** and GDP **(e)** is shown in dark blue.

**SFig. 4.**
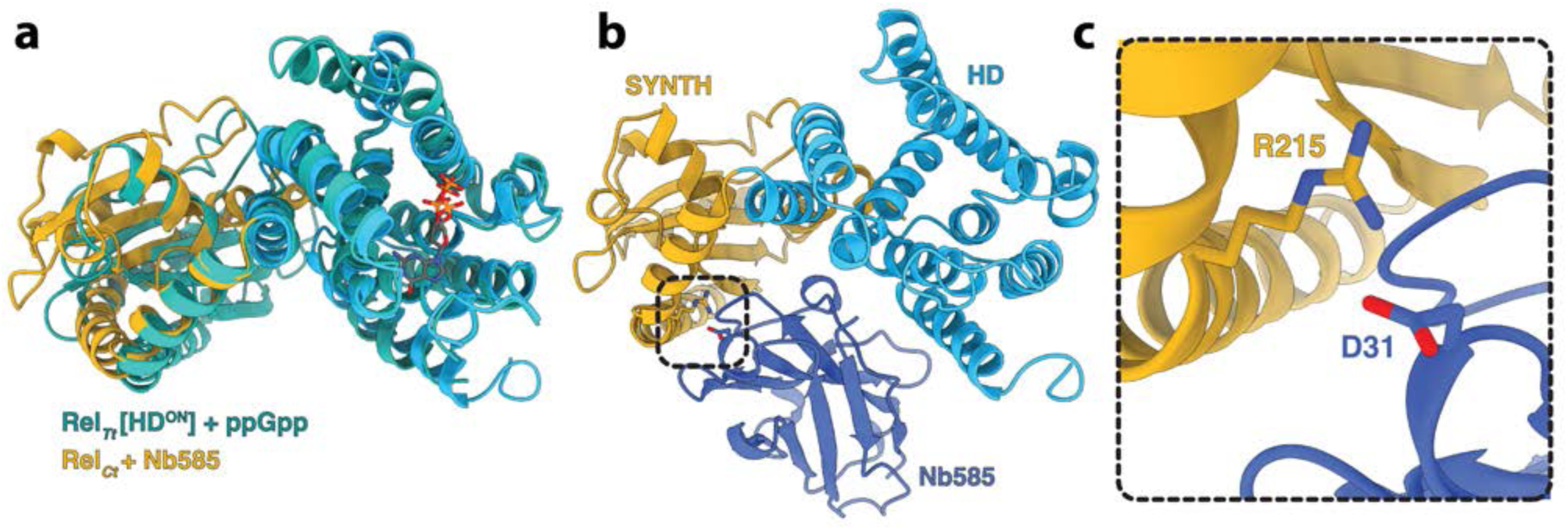
Nb585 stabilises a conformation that resembles the active HD state of Rel. **(a)** Structural superposition of Rel*_Ct_*^NTD^ in the closed conformation predicted in the complex with Nb585 onto the Rel*_Tt_*-ppGpp complex. The comparison supports the hypothesis that binding to Nb585 triggers a state that favours hydrolysis (illustrated by the presence of ppGpp in the HD site) and prevents synthesis (the SYNTH active site is largely occluded in this state). **(b)** Cartoon representation of the Rel*_Ct_*^NTD^-Nb585 complex coloured as in Fig. 5c. The main contact interface is highlighted by a black square. **(c)** Details of Rel*_Ct_*^NTD^-Nb585 complex interface highlighting key residues involved in binding including D31 of Nb585 (in blue) that forms a slat bridge with R215 of Rel*_Ct_*^NTD^ (in gold).

## SUPPLEMENTARY TABLES

**Supplementary Table 1.**
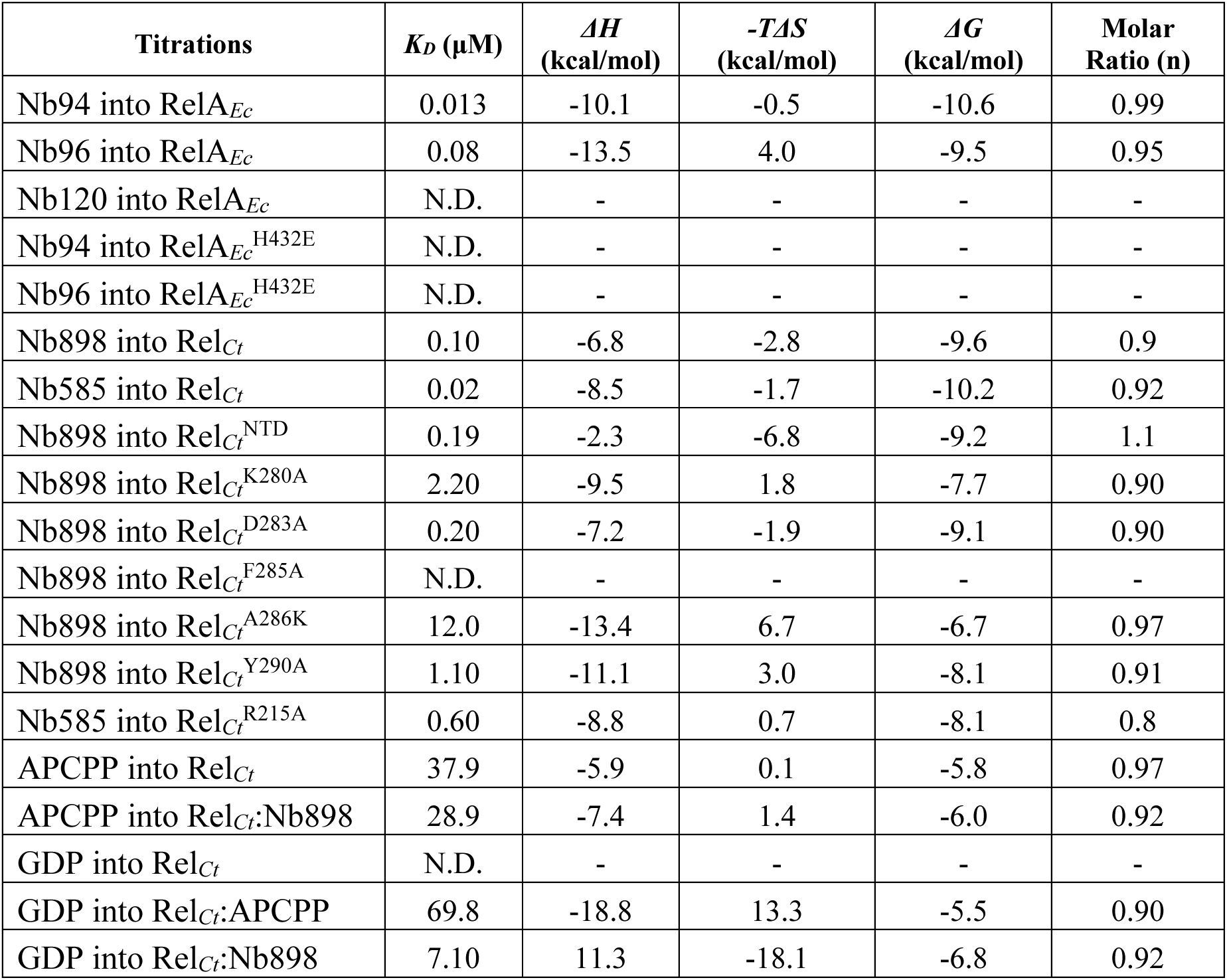
Binding parameters determined by Isothermal Titration Calorimetry (ITC). Experimentally determined binding thermodynamic parameters resulting from ITC measurements. N.D. stands for ‘not detectable’. All of the presented titrations are background-subtracted.

| Titrations | $K_D$ (μM) | $\Delta H$<br>(kcal/mol) | $-T\Delta S$<br>(kcal/mol) | $\Delta G$<br>(kcal/mol) | Molar<br>Ratio (n) |
| --- | --- | --- | --- | --- | --- |
| Nb94 into RelA<sub>Ec</sub> | 0.013 | -10.1 | -0.5 | -10.6 | 0.99 |
| Nb96 into RelA<sub>Ec</sub> | 0.08 | -13.5 | 4.0 | -9.5 | 0.95 |
| Nb120 into RelA<sub>Ec</sub> | N.D. | - | - | - | - |
| Nb94 into RelA<sub>Ec</sub><sup>H432E</sup> | N.D. | - | - | - | - |
| Nb96 into RelA<sub>Ec</sub><sup>H432E</sup> | N.D. | - | - | - | - |
| Nb898 into Rel<sub>Ct</sub> | 0.10 | -6.8 | -2.8 | -9.6 | 0.9 |
| Nb585 into Rel<sub>Ct</sub> | 0.02 | -8.5 | -1.7 | -10.2 | 0.92 |
| Nb898 into Rel<sub>Ct</sub><sup>NTD</sup> | 0.19 | -2.3 | -6.8 | -9.2 | 1.1 |
| Nb898 into Rel<sub>Ct</sub><sup>K280A</sup> | 2.20 | -9.5 | 1.8 | -7.7 | 0.90 |
| Nb898 into Rel<sub>Ct</sub><sup>D283A</sup> | 0.20 | -7.2 | -1.9 | -9.1 | 0.90 |
| Nb898 into Rel<sub>Ct</sub><sup>F285A</sup> | N.D. | - | - | - | - |
| Nb898 into Rel<sub>Ct</sub><sup>A286K</sup> | 12.0 | -13.4 | 6.7 | -6.7 | 0.97 |
| Nb898 into Rel<sub>Ct</sub><sup>Y290A</sup> | 1.10 | -11.1 | 3.0 | -8.1 | 0.91 |
| Nb585 into Rel<sub>Ct</sub><sup>R215A</sup> | 0.60 | -8.8 | 0.7 | -8.1 | 0.8 |
| APCPP into Rel<sub>Ct</sub> | 37.9 | -5.9 | 0.1 | -5.8 | 0.97 |
| APCPP into Rel<sub>Ct</sub>:Nb898 | 28.9 | -7.4 | 1.4 | -6.0 | 0.92 |
| GDP into Rel<sub>Ct</sub> | N.D. | - | - | - | - |
| GDP into Rel<sub>Ct</sub>:APCPP | 69.8 | -18.8 | 13.3 | -5.5 | 0.90 |
| GDP into Rel<sub>Ct</sub>:Nb898 | 7.10 | 11.3 | -18.1 | -6.8 | 0.92 |

**Supplementary Table 2.** X-ray data collection and processing. The *CC*_1/2_ criterion was used to determine the resolution range. Values for the outer shell are given in parentheses.

|  | **Rel<sub>CT</sub><sup>NTD</sup> : Nb898 complex** | **Rel<sub>CT</sub><sup>NTD</sup> : GDP : APCPP complex** |
| --- | --- | --- |
| Crystallization condition | 29 % w/v PEG 6000 0.2 M Sodium citrate | 27 % w/v PEG 3350 0.1 M Bis-Tris propane 0.2 M Lithium sulfate |
| Diffraction source | Soleil PX2 | Soleil PX2 |
| Wavelength (Å) | 0.9762 | 0.9801 |
| Temperature (K) | 100 | 100 |
| Detector | Dectris PILATUS 6M | Dectris Eiger |
| Detector distance (mm) | 602.7 | 342.5 |
| Space group | C2 | P2<sub>1</sub> |
| *a*, *b*, *c* (Å) | 196.56 96.47 62.98 | 92.63 92.31 122.29 |
| $\alpha$, $\beta$, $\gamma$ (°) | 90.00 102.16 90.00 | 90.00 94.80 90.00 |
| Resolution range (Å) | 96.07 - 3.15 (3.31 – 3.15) | 50.85 - 2.65 (2.89 - 2.65) |
| Total No. of reflections | 76808 (3781) | 248789 (11760) |
| No. of unique reflections | 11630 (581) | 46040 (2302) |
| Completeness (%) | 88.8 (89.8) | 93.5 (53.5) |
| Redundancy | 6.7 (6.1) | 5.4 (5.1) |
| $\langle I/\sigma(I) \rangle$ | 5.1 (1.1) | 9.0 (1.1) |
| *R*<sub>merge</sub> | 0.619 (3.235) | 0.15 (1.33) |
| *CC*<sub>1/2</sub> | 0.956 (0.122) | 0.994 (0.411) |
| Overall *B* factor / Wilson plot (Å<sup>2</sup>) | 48.2 | 63.8 |
| *R*-factor (%) | 24.1 | 21.6 |
| *R*<sub>free</sub>-factor (%) | 28.3 | 23.4 |
| Ramachandran profile (%) |  |  |
| Core | 95.4 | 98.3 |
| Allowed | 4.6 | 1.7 |
| Outliers | 0.0 | 0.0 |
| R.m.s. deviations |  |  |
| Bond lengths (Å) | 0.005 | 0.013 |
| Bond angles (°) | 1.084 | 1.63 |
| Number of atoms | 6404 | 11095 |
| Macromolecules | 6397 | 10679 |
| Solvent | 4 | 201 |
| Other | 3 | 215 |
| B-factors (Å<sup>2</sup>) |  |  |
| All atoms | 40.6 | 70.6 |
| Macromolecules | 40.6 | 70.5 |
| Solvent atoms | 25.1 | 57.0 |
| Other atoms | 56.7 | 64.2 |
| PDB ID | 9S6K | 9S6L |

## Online Methods

### Multiple sequence alignment

Sequences were aligned with MAFFT v7.164b with the L-INS-i strategy^54^ and visualised with Jalview^55^.

### Enzymatic assays

Purified protein was incubated in HEPES Polymix buffer (as in Takada et. al., 2020; Tamman et. al., 2020; Takada et. al., 2021; Roghanian et. al., 2021) with HEPES pH 7.5 and 0.5 mM TCEP to a final concentration of 0,25 µM. The Nbs (2.5 µM) or 70S ribosomal complexes (1.5 µM) (*E. coli*) were added prior to a 5 min incubation at room temperature. In case of deacylated tRNA (1 µM) (mix) and pppGpp-mediated (100 µM) stimulation of (p)ppGpp synthesis, these compounds were added after incubation with 70S ribosomal complexes for 5 minutes and incubated for a subsequent 5 minutes at room temperature. When monitoring the ppGpp-synthesis reaction, GDP was added to 1 mM final concentration to the mixture before incubation for 1 min at 40 °C. Synthesis of ppGpp was initiated by the addition of ATP (at 40 °C) to a final concentration of 1 mM. When monitoring the ppGpp-hydrolysis reaction, the reaction mix was incubated at 40 °C for 1 minute before adding the substrate (ppGpp) at a final concentration of 1 mM. The reaction mixtures were incubated in a mixing heat block at 40°C under rigorous shaking at 600 r.p.m. Quenching of the enzymatic reactions was achieved by the addition of formic acid (10 %). The reaction mixes were stored on ice and centrifuged for 1 minute at 4 °C before analysis by means of HPLC.

HPLC was performed on a Shimadzu i-series device using an ACE-equivalence C18 HPLC column (3 µM, 30 x4.6 mm, Avantor). Nucleotides were separated by reverse phase applying a gradient (0-100 %) of buffer B (100 mM K phosphate pH 6.4, 5 mM tetrabutyl ammonium bromide (TBAB), 15 % acetonitrile) to buffer A (10 mM K phosphate buffer (pH 6.4, 5 mM TBAB, 0.1 % acetonitrile) at a flowrate of 0.8 mL / minute for 10 minutes. The nucleotides were monitored by absorbance at 254 nm and concentration of GDP and ppGpp was derived from calibration curves using the surface area of the respective peaks. All measurements were analysed using the Labanalysis software. Turnovers were obtained from linear regression of the GDP or ppGpp synthetized in time. All experiments were performed in triplicates.

### Growth assays on SMG (Serine, Methionine and Glycine) plates

The SMG plate assay was performed as described earlier^17^. The strains were grown overnight in liquid M63^44^-glucose minimal medium supplemented with 0.4% glucose, 5 μg/mL thiamine, 100 μg/mL each of L-Arginine, L-Histidine, L-Leucine, L-Threonine and 100 µg/mL ampicillin. Optionally, 0.5 mM IPTG was used to induce nanobody expression and 1 mM each of L-Serine, L-Methionine, and L-Glycine were added to repress growth of RelA-deficient strains. The cultures were adjusted to OD_600_ 0.5 and 10 fold dilutions were plated on solid M63, SMG and SMG-IPTG plates^43^ supplemented with 100 µg/mL ampicillin.

### Virulence assays in *G. mellonella*

For inoculation, *G. mellonella* larvae were incubated 1h at 4 °C before injection. Overnight culture of *E. coli* Ec156^45^ transformed with the vector expressing the nanobodies were washed and diluted to a cell density equivalent to 100 CFU in 0.9 % NaCl, 10 µL and were injected in the bottom left proleg of each larva. Twenty larvae were inoculated per strain and incubated at 37 °C in the dark. Viability of the larvae was scored every 12 h.

### Protein purification

Overexpression and purification of *E. coli* RelA and *C. tepidum* Rel was performed as described earlier^19,56^. To purify Rel*_Ct_*^NTD^ and variants, clarified cell lysate of N-terminally His-TEV-tagged proteins was loaded onto a HisTrap HP 1 mL column (GE healthcare) pre-equilibrated with 5 column volumes of the binding buffer (50 mM Hepes pH 7.5, 500 mM KCl, 500 mM NaCl, 10 mM MgCl_2_, 1 mM TCEP and 0.002 % mellitic acid) 1 pastil of protease inhibitors cocktail (Roche)), respectively. A washing step with binding buffer preceded elution of the protein by a linear gradient of 0-500 mM imidazole. Elution fractions were pooled for size exclusion chromatography with a HiLoad 16/600 Superdex 200 PG column (GE healthcare) pre-equilibrated with respective binding buffer. Purified truncates were treated with TEV-protease in a 1:100 molar ratio, overnight, at room temperature to remove the His-tag. The protease and cleaved tag were removed by reverse IMAC using a TALON gravity flow column followed by SEC to the protein buffer. The proteins were concentrated to appropriate concentration using ultrafiltration units with 30 000 Da cut-off (Amicon Ultra, Merck Milipore). Purity of the samples’ preparation was assessed spectrophotometrically and by SDS-PAGE.

In all cases nanobodies were produced in *E. coli* WK6 cells^39^ transformed with pMesy4 plasmid carrying the Nb-genes. Cells were grown in TB broth and expression was induced with 500 μM IPTG overnight at 28 °C. Periplasmatic extraction of the nanobodies was done by osmotic shock was achieved as previously described^41^. The His-tagged Nbs were purified using a His-trap Ni-NTA IMAC-column (Cytiva) on an AKTA pure FPLC-system at 8 °C in 25 mM HEPES 7.5; 250 mM NaCl; 1 mM TCEP. Elution of the was performed by applying a gradient of imidazole (0-500 mM). IMAC was followed by buffer exchange on SEC column (Superdex75 increase 10/300 GL, Cytiva) to HEPES 7.5; 250 mM NaCl; 1 mM TCEP.

### Isothermal affinity calorimetry (ITC)

All titrations were performed with an Affinity-ITC (TA instruments) at 15 °C using a constant titration volume of 2μL. Samples were incubated and degassed for 10 minutes prior to the ITC measurement. Nucleotide stocks of Adenosine-5’-[(α,β)-methyleno]triphosphate (ApCpp), ppGpp, pppGpp (Jena Biosciences) and GDP (Sigma Aldrich) of 650–670 mM were diluted in 50 mM HEPES pH 7.5; 500 mM KCl; 500 mM; NaCl; 10 mM MgCl_2_; 1 mM TCEP; 0.002 % mellitic acid,to a 16-22 fold excess relative to the protein in the ITC cell. The purified RSH-enzymes were concentrated by ultrafiltration (Amicon Ultra, 0.5 ml, 30 kDa, Merck Millipore) at 3,000g at 15 °C to a final concentration of 44–1800 μM. All ITC measurements were performed using a constant stirring rate of 75 r.p.m. All data were processed and analyzed using the NanoAnalyse and Origin software packages, the data is summarized in **Supplementary Table S1**.

### Crystallisation of Rel_*Ct*_^NTD^/Nb898-complex

Crystals of Rel*_Ct_*^NTD^ in complex with GDP and APCPP were obtained by incubating Rel*_Ct_*^NTD^ at 10 mg/mL with 10 mM (final concentration) of the nucleotides. Crystals were obtained in F8 condition of the LMB crystallization screen (Hampton research) at 20 °C using the sitting-drop vapour-diffusion technique with Swissci (MRC) 96-well 2-drop UVP sitting-drop plates. The 200 nL drops of 1:1 crystallization solution/protein sample, were deposited under humidity-controlled conditions using the Mosquito robotic system (TTP Labtech). The drops were equilibrated to 80 μL of the crystallization solution on the plate.

To obtain crystals of Rel*_Ct_*^NTD^ in complex with Nb898, the nanobody was added in a 5-fold molar excess to purified Rel*_Ct_*^NTD^ and incubated for 5 minutes at room temperature. SEC on an Åkta pure FPLC system at 8 °C with a Superdex 200 increase 10/300 GL column (cytiva) equilibrated with the protein buffer at 2% glycerol was performed on the mixture to isolate the protein complex. Formation of the complex was verified on denaturing SDS-PAGE. The protein complex was concentrated by ultrafiltration using a 3000 Da cut-off (Amicon Ultra, Merck Milipore) to 14 mg/mL as estimated by OD_280nm_ (on a NanoDrop microvolume spectrophotometer, Thermo-Fisher Scientific) using theoretical extinction coefficients. Crystals were obtained in D9 condition of the LMB crystallisation screen (Hampton research) at 20 °C using the sitting-drop vapour-diffusion technique with Swissci (MRC) 96-well 2-drop UVP sitting-drop plates. The 200 nL drops of 1:1 crystallization solution/protein sample, were deposited under humidity-controlled conditions using the Mosquito robotic system (TTP Labtech). The drops were equilibrated to 80 μL of the crystallization solution on the plate.

### Structure determination

X-ray diffraction data of crystals of Rel*_Ct_*^NTD^:Nb898 were collected at the PX2 beamline of the Soleil synchrotron (Gif-sur-Yvette, France) on an Eiger detector. Prior to data collection, crystals the crystals were transferred to a suitable cryoprotectant solution containing 38% of 2-Methyl-1,3-propanediol (MPD) and flash-cooled in liquid nitrogen. The data were processed using XDS suite^37^ and scaled with Aimless. The anisotropic analysis of the diffraction data suggested a resolution of 3.15 Å (with diffraction limits of 3.39 Å in a*, 4.99 Å in b* and 2.73 Å in c*). The structure was solved by molecular replacement performed with Phaser^38^ using the coordinates of nanobody T4^39^ (PDB ID: 6GJS) after removing the CDR regions and the coordinates of Rel*_Tt_*^NTD^. Initial automated model building was performed with Buccaneer^42^ which partially completed the Nb898 and further improved with the MR-Rosetta suite from the Phenix package^43^. After several iterations of manual building with Coot^41^ and maximum likelihood refinement as implemented in Buster/TNT^44^, the model was refined to R/Rfree of 0.243/0.283.

In the case of the Rel*_Ct_*^NTD^:GDP:APCPP complex X-ray diffraction data were collected at the PX2 beamline of the Soleil synchrotron (Gif-sur-Yvette, France) on an Eiger detector. Crystals were cryo-protected in 20%glycerol and 2% DMSO and flash-cooled in liquid nitrogen prior to data collection. The data were processed using XDS suite^37^ and scaled with Aimless. Anisotropic analysis of the diffraction data suggested a resolution of 2.65 Å (with diffraction limits of 3.07 Å in a*, 2.65 Å in b* and 2.80 Å in c*). The structure was solved by molecular replacement performed with Phaser^38^ using the coordinates of the separated HD and SYNTH domains of Rel*_Ct_*^NTD^ (from the Rel*_Ct_*^NTD^-Nb898 complex). Automated model building was performed MR-Rosetta suite from the Phenix package^43^. After several iterations of manual building with Coot^41^ and maximum likelihood refinement as implemented in Buster/TNT^44^, the model was refined to R/Rfree of 0.187/0.228. The X-ray data collection and refinement statistics of both datasets are summarized in **Supplementary Table S2**.

### Hydrogen-deuterium exchange mass spectrometry

HDX-MS experiments were performed on an HDX platform composed of a Synapt G2-Si mass spectrometer (Waters Corporation) connected to a nanoACQUITY ultraperformance liquid chromatography (UPLC) system, as described^57^. Samples of Rel*_Ct_*^NTD^, Nb898, Nb585 the Rel*_Ct_*^NTD^:Nb898 and Rel*_Ct_*^NTD^:Nb585 complexes were prepared at a concentration of 50 to 70 μM. For each experiment, 5 μl of sample was incubated for 0.5, 5, or 60 min in 95 μl of labeling buffer L [50 mM HEPES, 500 mM KCl, 500 mM NaCl, 2 mM MgCl_2_, 1 mM TCEP, and 0.002% mellitic acid (pH 7.5)] at 20 °C. The nondeuterated reference points were prepared by replacing buffer L by equilibration buffer E [50 mM Hepes, 500 mM KCl, 500 mM NaCl, 2 mM MgCl_2_, 1 mM TCEP, and 0.002% mellitic acid (pH 7.5)]. After labeling, the samples are quenched by mixing with 100 μl of prechilled quench buffer Q [1.2% formic acid (pH 2.4)]. Seventy microliters of the quenched samples were directly transferred to the Enzymate BEH Pepsin Column (Waters Corporation) at 200 μl/min and at 20 °C with a pressure of 8.5 kpsi. Peptic peptides were trapped for 3 min on an ACQUITY UPLC BEH C18 VanGuard precolumn (Waters Corporation) at a flow rate of 200 μl/min in water [0.1% formic acid in HPLC water (pH 2.5)] before being eluted to an ACQUITY UPLC BEH C18 column for chromatographic separation. Separation was performed with a linear gradient buffer (7 to 40% gradient of 0.1% formic acid in acetonitrile) at a flow rate of 40 μl/min. Identification of peptides and deuteration uptake analysis was performed on the Synapt G2Si in (ESI+) - HDMSE mode (Waters Corporation). Leucine enkephalin was applied for mass accuracy correction, and sodium iodide was used as calibration for the mass spectrometer. HDMSE data were collected by a 20- to 30-V transfer collision energy ramp. The pepsin column was washed between injections using pepsin wash buffer [1.5 M Gu-HCl, 4% (v/v) methanol, and 0.8% (v/v) formic acid]. A blank run was performed between each sample to prevent significant peptide carryover. Optimized peptide identification and peptide coverage for all samples were performed from undeuterated controls (five replicates). All deuterium time points were performed in triplicate.

